# Cholesterol, PtdIns(4,5)P_2_, and Actin Regulate BK Channel Nanoscale Organization

**DOI:** 10.64898/2026.04.20.719652

**Authors:** Roya Pournejati, Michael Ma, Jessica M. Huang, Lizbeth de la Cruz, Claudia M. Moreno, Oscar Vivas

## Abstract

BK channels form nanoscale clusters, but the membrane and cytoskeletal mechanisms that organize these structures remain poorly understood. Using super-resolution imaging and FRAP, we dissect how cholesterol, PtdIns(4,5)P_2_, and actin govern BK channel organization expressed heterologously in tsA-201 cells and expressed endogenously in INS-1 cells. Cholesterol depletion produced opposite outcomes across these systems, enlarging clusters in heterologous cells but reducing both size and density in INS-1 cells. PtdIns(4,5)P_2_ displayed a biphasic regulatory profile: both enrichment and depletion enlarged clusters, revealing that BK assemblies are tuned to a narrow PtdIns(4,5)P_2_ set point. Actin disruption consistently reduced cluster size and density. Together, these findings show that cholesterol, PtdIns(4,5)P_2_, and actin are not only functional regulators of BK channels but also key determinants of their nanoscale organization. Across perturbations, changes in channel density strongly predicted changes in cluster size, identifying density as a central organizing principle.

**Significance statement:** This study shows that the nanoscale organization of BK channels is dynamically controlled by membrane lipids and the cytoskeleton, demonstrating that ion channel organization is not fixed but actively shaped by the local membrane environment. Across perturbations, cluster size scales with channel density, underscoring density as a major driver of BK nanoscale organization.

## Introduction

Ion channels are integral membrane proteins that operate within complex lipid–protein environments. Channel activity is not determined solely by intrinsic gating mechanisms but is strongly influenced by the composition and organization of the surrounding membrane [1–3]. In addition to providing structural integrity to the membrane, lipids and cytoskeletal elements also support and regulate channel conductance and responsiveness to physiological signals [4]. These channel-membrane interactions are increasingly recognized as critical determinants of excitability in diverse cell types [5]. However, the role of membrane composition in organizing ion channels at the nanoscale, including their clustering, distribution, and mobility, has remained largely unexplored. Addressing this gap is important, as nanoscale organization has been proposed to play a role in ion channel and broader protein function.

BK (large-conductance, calcium- and voltage-activated potassium) channels are particularly sensitive to the membrane environment. These channels play essential roles in controlling neuronal firing, vascular tone, and endocrine secretion [6]. Previous studies have shown that cholesterol [7], phosphatidylinositol 4,5-bisphosphate (PtdIns(4,5)*P*_2_) [8], and the actin cytoskeleton [9] alter BK channel activity. While functional modulation is well established, no studies have directly examined how membrane composition influences the clustering and spatial organization of BK channels.

To address this gap, we focused on cholesterol, PtdIns(4,5)*P*_2_, and actin, three membrane associated components with well-established molecular interfaces to BK channels. Cholesterol interacts with cholesterol recognition amino acid (CRAC) motifs within transmembrane domains, influencing gating and lipid sensitivity [10]. PtdIns(4,5)*P*_2_ engages clusters of positively charged residues in the cytoplasmic domain, where electrostatic interactions stabilize channel open states [11]. Actin interacts through indirect scaffolding connections and through direct cortactin-mediated binding motifs that anchor BK channels to the cortical cytoskeleton [9].

In this study, we combined high-resolution imaging with quantitative cluster analysis to assess BK channel distribution expressed both heterologously in tsA-201 cells and endogenously in INS-1 cells, allowing for comparison between cellular contexts. In tsA-201 cells, we further applied fluorescence recovery after photobleaching (FRAP) to measure BK channel mobility, providing complementary insight into dynamic behavior. Our findings reveal that cholesterol exerts cell-type dependent effects, whereas PtdIns(4,5)*P*_2_ consistently promotes clustering, and actin provides structural stabilization. Together, these results establish membrane composition and cytoskeletal integrity as key determinants of BK channel organization and dynamics.

## Material and methods

### Cell culture and transfection

INS-1 pancreatic β-cells and tsA-201 cells were used to investigate BK channel clustering in both endogenous and heterologous contexts. INS-1 cells were cultured in RPMI 1640 medium (Gibco) supplemented with 10% fetal bovine serum, 1 mM sodium pyruvate, 10 mM HEPES, 50 μm β-mercaptoethanol and 0.2% penicillin/streptomycin. Cultures were maintained at 37 °C in a humidified 5% CO_2_ atmosphere and passaged weekly. For experiments involving phosphoinositide modulation, INS-1 cells were transfected with DNA constructs encoding PIP5K and PIP5P, using Lipofectamine 3000 (Invitrogen; RRID: L30000), and used for live imaging or immunofluorescence 24 hours post-transfection.

tsA-201 cells were cultured in Dulbecco’s Modified Eagle Medium (DMEM; Gibco), supplemented with 10% fetal bovine serum and 0.2% penicillin/streptomycin, and maintained at 37 °C in a 5% CO_2_ atmosphere. Cultures were passaged twice a week. For heterologous expression of BK channels, tsA-201 cells were plated and transfected with 0.1 μg of DNA per plasmid using Lipofectamine 3000. After 4 hours of transfection, cells were seeded onto poly-D-lysine–coated glass coverslips and incubated for 24 hours prior to live imaging or immunofluorescence assays.

### DNA construct and plasmids

DNA clones of BK channels, PH-Place GFP, and PIP5K were obtained from Addgene (RRID: SCR_002037). PIP5P and D4H-mCherry were generously provided by Bertil Hille. BK-GFP was a kind gift from James Trimmer (University of California, Davis, CA). No BK channel auxiliary subunits were transfected.

### Antibodies

BK channels were detected using the anti-Slo1 mouse monoclonal antibody clone L6/60 or Anti-Maxi K/BK channel rabbit monoclonal antibody clone 3D15. Donkey anti-mouse Alexa-647 and donkey anti-rabbit Alexa-555 were used as secondary antibodies. All were purchased from Molecular Probes. Antibody specificity was tested in untransfected tsA-201 cells.

### Treatments to modulate lipid levels and actin

To modulate phosphoinositide signaling, cells were transfected with DNA constructs encoding PIP5K and PIP5P and incubated for 24 hours prior to imaging or functional assays. To assess the effect of rapid and substantial depletion of PtdIns(4,5)*P*_2_, cells were transfected with muscarinic acetylcholine receptor type 1 (M1R). The M1R signaling pathway was activated with 10 μM oxotremorine M, and the resynthesis was inhibited with 200 nM GSK-A1 in some experiments. To assess the impact of cholesterol and cytoskeletal integrity, cells were treated with either 5 mM methyl-β-cyclodextrin (MβCD) for cholesterol depletion or 5 μM cytochalasin D (Cyt D) to disrupt the actin cytoskeleton. Both treatments were performed in Ringer’s solution at 37 °C. MβCD was applied for 60 minutes, while Cyt D was applied for 30 minutes. Following each treatment, cells were washed three times with fresh Ringer’s solution every 5 minutes. For visualization of the actin cytoskeleton, live cells were labeled with CellMask™ Actin stain according to the manufacturer’s protocol. Ringer’s solution was prepared with the following components (in mM): 155 NaCl, 4.5 KCl, 1 MgCl_2_, 2 CaCl_2_, 10 HEPES, and 10 glucose. The pH was adjusted to 7.4 using NaOH, and osmolarity was maintained at ∼320 mOsm. All treatments and imaging procedures were conducted using freshly prepared solutions. To evaluate the effects of PtdIns(4,5)*P*_2_ and cholesterol modulation, cells were transfected with PH-PLCδ-GFP or D4H-mCherry, which visualize PtdIns(4,5)*P*_2_ and cholesterol, respectively. For each cell, mean fluorescence intensities for the cell border and interior were measured. These mean fluorescence intensities were used to calculate each cell’s plasma membrane-to-cytoplasmic fluorescence intensity ratio, which was then used to compare treatment conditions.

### Immunostaining

Cells were fixed for 10 minutes at room temperature using freshly prepared 4% paraformaldehyde. To quench residual aldehydes, samples were treated with 0.1% sodium borohydride for 5 minutes, followed by multiple washes with phosphate-buffered saline (PBS). To reduce nonspecific antibody binding, cells were blocked with 3% bovine serum albumin (BSA; Thermo Scientific) in PBS. Cells were permeabilized with 0.25% Triton X-100 in PBS for 1 hour at room temperature. Primary antibodies were diluted to 10 μg/ml in blocking solution and applied overnight at 4 °C. After thorough washing, secondary antibodies were added at a concentration of 2 μg/ml and incubated for 1 hour at room temperature. All wash steps consisted of three PBS rinses, each followed by gentle agitation for 5 minutes.

### High-resolution imaging

Confocal imaging was performed using a Zeiss LSM 880 inverted microscope equipped with Airyscan detection and operated via ZEN Black v2.3 software. A Plan-Apochromat 63x oil immersion objective (numerical aperture 1.4) was used for all acquisitions. Fluorescent signals were excited using a combination of laser lines: 405 nm diode, 458–514 nm argon, 561 nm, and 633 nm. Emission was collected through appropriate filter sets using a 32-channel GaAsP Airyscan detector. Cells were imaged in Ringer’s solution. Image processing and quantification were performed using ImageJ (NIH).

### Super-resolution imaging

To visualize BK channels in tsA-201 and INS-1 cells, we performed direct stochastic optical reconstruction microscopy (dSTORM) using the ONI Nanoimager system. The microscope was equipped with a 100x oil immersion objective (numerical aperture 1.4), an XYZ closed-loop piezoelectric stage (model 736), and three emission channels split at 488, 555, and 640 nm. Imaging was conducted at 35 °C. Fixed and immunostained cells were immersed in GLOX imaging buffer containing 10 mM β-mercaptoethylamine (MEA), 0.56 mg/ml glucose oxidase, 34 μg/ml catalase, and 10% (w/v) glucose in Tris-HCl. Single-molecule localization data were acquired and initially processed using NImOS software (ONI, version 1.18.3). Localization maps were exported as TIFF files with a pixel resolution of 5 nm. Localization maps were further processed in ImageJ (NIH), where maps were thresholded and binarized to isolate labeled structures. Particles smaller than 400 nm^2^ were excluded to account for the resolution limits of dSTORM (∼20 nm) and the estimated footprint of BK channels.

### Fluorescence Recovery After Photobleaching (FRAP)

BK-GFP fluorescence was excited using a 488 nm argon laser. Imaging was performed at the cell footprint, which was identified by a uniform fluorescence intensity across the cell edge and interior, indicating membrane localization without cytoplasmic interference. Bleaching was performed by drawing a 3.5 μm wide circular region of interest and then simultaneously restricting the excitation laser to that area while increasing its power to the maximum. A bleaching pixel dwell of 9.31 µs for 8 iterations was used to ensure significant photobleaching while minimizing BK channel diffusion outside the region during bleaching. The region of interest was selected to purposely exclude distinctive bright puncta larger than 50,000 nm^2^. Fluorescence was monitored over 80 seconds, and reported as intensity values normalized to prebleach fluorescence. The half-time of recovery and bleach radius were used to calculate channel diffusion, as described by Kang et al [12]. The mobile fraction was calculated in a two-step process. The nadir of the fluorescence trace, defined as the time of bleaching, was first set to 0, shifting the entire trace downward. Then, the ratio was calculated between the average values of the final 50 frames and all frames before bleaching.

### Statistical analysis and data representation

Quantitative data were analyzed using Microsoft Excel and GraphPad Prism. Image processing and quantification were performed in ImageJ (NIH). Statistical comparisons were conducted using one-way ANOVA for parametric datasets and the Mann–Whitney U test for non-parametric distributions. A p-value threshold of less than 0.05 was considered indicative of statistical significance. The number of cells included in each experimental condition is specified in the corresponding figure legends. Data in bar charts are shown as mean ± SEM. Data in FRAP curves and cumulative frequency distributions are shown as individual, fainter curves and a bolded average curve.

## Results

### Cholesterol depletion promotes larger clusters and reduces BK channel mobility

To investigate how membrane composition influences BK channel organization and dynamics, we first examined the effects of cholesterol depletion in tsA-201 cells expressing D4H-mCherry, a cholesterol sensor. Cells were treated with 5 mM methyl-β-cyclodextrin (MβCD) to extract membrane cholesterol, and cholesterol distribution was visualized using D4H-mCherry. Quantitative analysis revealed a 17-fold reduction in plasma membrane localization of D4H-mCherry following MβCD treatment, confirming effective cholesterol depletion (Figure 1A–B).

**Figure 1.**
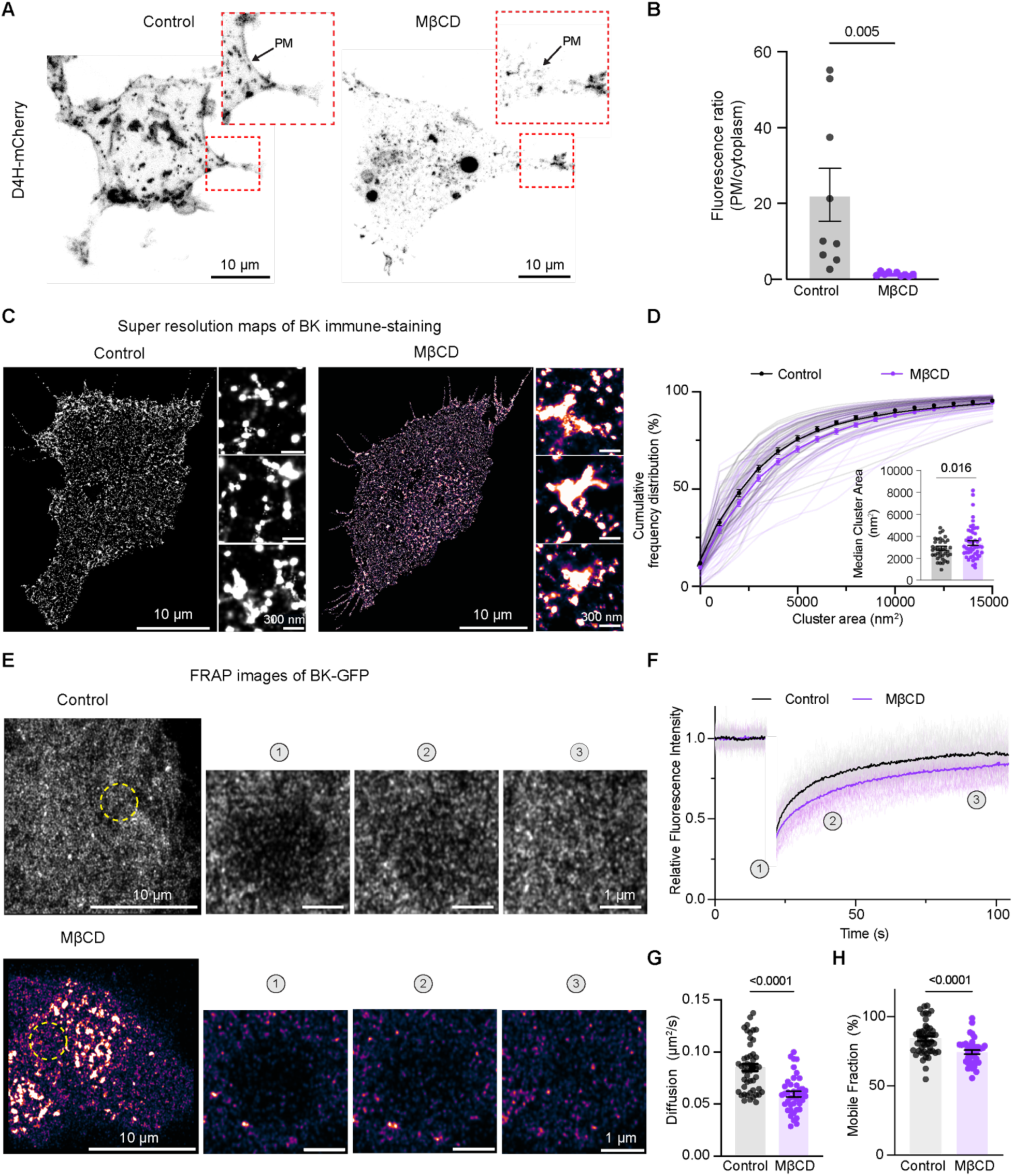
Cholesterol depletion reduces BK channel mobility and promotes larger clusters. **A.** Representative confocal images of tsA-201 cells transfected with D4H-mCherry. Left: control; right: 5 mM MβCD treatment. Scale bar: 10 µm. **B.** Comparison of D4H-mCherry fluorescence intensity ratio between control (n=9) and MβCD-treated (n=10). **C.** Representative super-resolution localization maps of BK immunostaining. Left: control; Right: treated. Scale bars: 10 µm and 300 nm in magnification. **D.** Cumulative frequency plot comparing BK cluster size distributions between control (n=38) and MβCD-treated (n=53). Statistical significance was tested using a 2-way ANOVA. P-value was 0.016. Inset: scatter plot showing median cluster size in nm². Statistical significance was tested using a two-tailed t-test. **E.** Representative FRAP time-lapse images of BK-GFP. Top: control; Bottom: MβCD. Scale bar: 10 µm and 1 µm in magnification. **F.** FRAP recovery curves comparing BK-GFP mobility. **G.** Scatter dot plot comparing BK channel diffusion between control and MβCD-treated. **H**. Scatter dot plot comparing mobile fraction of BK clusters between control and MβCD-treated. In panels E to H, data come from n= 49 in control and n = 42 in MβCD. Statistical significance in all scatter plots was tested using a two-tail t-test.

We assessed the spatial organization of BK channels using direct stochastic optical reconstruction microscopy (dSTORM). Localization maps of BK immunostaining (Figure 1C), rendered at 5 nm pixel resolution, revealed multi-pixel aggregates consistent with BK channel clusters. Interpretation of these maps requires consideration of methodological limitations. Each functional BK channel is a tetramer composed of four α-subunits [13], and in principle, each subunit can bind a single primary antibody. dSTORM cannot distinguish whether signals arise from labeling of one or all four α subunits within a single tetramer or from labeling only one α subunit within multiple tetramer complexes. To address this ambiguity, we evaluated two scenarios. In the first scenario, we assumed that all four α-subunits of the BK channel tetramer are labeled. Cryo-EM structures indicate that a BK tetramer spans approximately 13 × 13 nm (≈169 nm²) [14]. Incorporation of two antibody layers (primary and secondary) would expand the footprint by about 14 nm in each dimension, yielding an estimated area of ∼41 × 41 nm (≈1681 nm²). In this case, particles smaller than ∼1681 nm² likely represent individual tetramers, whereas larger particles correspond to clusters of multiple tetramers.

In the second scenario, only a single antibody binds per tetramer. This stoichiometry may result from steric constraints at the S9–S10 segment, which could restrict labeling of additional α subunits within the tetramer. To distinguish between these scenarios, we used two antibodies against the BK C-terminal domain, raised in different species and labeled with distinct secondary antibodies conjugated to separate fluorophores. Super-resolution dSTORM imaging revealed colocalization in only 12.2 ± 0.3% of particles (Figure S1-1A), indicating that about 90% of tetramers bind just a single antibody. Furthermore, BK channel density (∼5 clusters per μm², Figure S1-1B) and median cluster area (∼3600 nm², Figure S1-B) were similar for both antibodies.

Based on these results, we favor the second scenario, in which steric constraints limit antibody binding to a single site per tetramer. Therefore, we consider ∼169 nm² as the footprint of a single BK channel. However, we increased the cutoff for a single channel to 400 nm², since the detection accuracy of the technique is 20 nm, not 13 nm, in both the X and Y axes.

Having established the properties of single-channel detections, we next evaluated how cholesterol depletion modifies BK cluster organization. Representative d-STORM maps of BK channel clusters in control and MβCD-treated cells are shown in Figure 1C. MβCD shifted the cumulative frequency distribution toward larger cluster sizes (Figure 1D; 2-way ANOVA between cumulative curves, *p = 0.016*). Consistent with this shift, the median cluster size increased from 2801 ± 147 nm² in control cells to 3429 ± 208 nm² after MβCD treatment (p = 0.016), representing an increase of around 630 nm².

To evaluate the impact of cholesterol depletion on BK channel mobility, we performed fluorescence recovery after photobleaching (FRAP). Representative time-lapse images (Figure 1E) captured fluorescence recovery over time in control and MβCD-treated cells. Recovery curves (Figure 1F) revealed a marked reduction in recovery amplitude in the MβCD condition. Importantly, the diffusion coefficient of BK-GFP decreased from 0.085 ± 0.003 μm²/s in control cells to 0.060 ± 0.003 μm²/s following MβCD treatment (Figure 1G), indicating a 30% reduction in lateral mobility of the mobile population. Consistent with this, the mobile fraction was significantly reduced from 84 ± 2 in control to 74 ± 1% after cholesterol depletion (Figure 1H). As a control for the mobile fraction in bleached regions, we measured the percentage of fluorescence change in unbleached regions (Figure S1-2) and found no decrease in fluorescence in unbleached regions. These results suggest that MβCD limits the proportion of BK clusters capable of dynamic exchange within the membrane.

### PtdIns(4,5)*P*_2_ enrichment promotes larger clusters whereas acute depletion increases BK channel mobility

To modulate plasma membrane PtdIns(4,5)P_2_ levels, we used complementary enzymatic and receptor-mediated approaches. Overexpression of PIP5K or PIP5P selectively increased or decreased PtdIns(4,5)P_2_ by phosphorylating PI(4)P to PtdIns(4,5)P_2_ or dephosphorylating PtdIns(4,5)P_2_ to PI(4)P, respectively (Figure S2-1). Substantial and fast depletion was achieved by activating overexpressed M1 muscarinic receptors (M1R) with Oxotremorine-M (Oxo-M, supplementary video S2-1), which triggers PLCβ-dependent hydrolysis of PtdIns(4,5)P_2_ [15]. A schematic of the strategy and biochemical pathway is shown in Figure S2-1.

We validated these manipulations using the PtdIns(4,5)P_2_ biosensor PH-PLCδ-GFP. PIP5K expression produced a clear increase in membrane-associated fluorescence, and quantification of the ratio of the fluorescence at the plasma membrane and the fluorescence in the cytosol confirmed a 36% elevation relative to control (Figure 2A–B). With this control established, we could examine how altered PtdIns(4,5)P_2_ abundance affects BK channel organization using dSTORM. Under control conditions, BK channels formed discrete clusters of variable size and density. PtdIns(4,5)P_2_ enrichment shifted the cumulative frequency distribution toward larger cluster sizes (Figure 2D; 2-way ANOVA between cumulative curves, p = 0.001). Consistent with this shift, the median cluster size increased from 2661 ± 178 nm² in control cells to 3747 ± 349 nm² after PIP5K overexpression (p = 0.008), representing an increase of 1086 nm².

**Figure 2.**
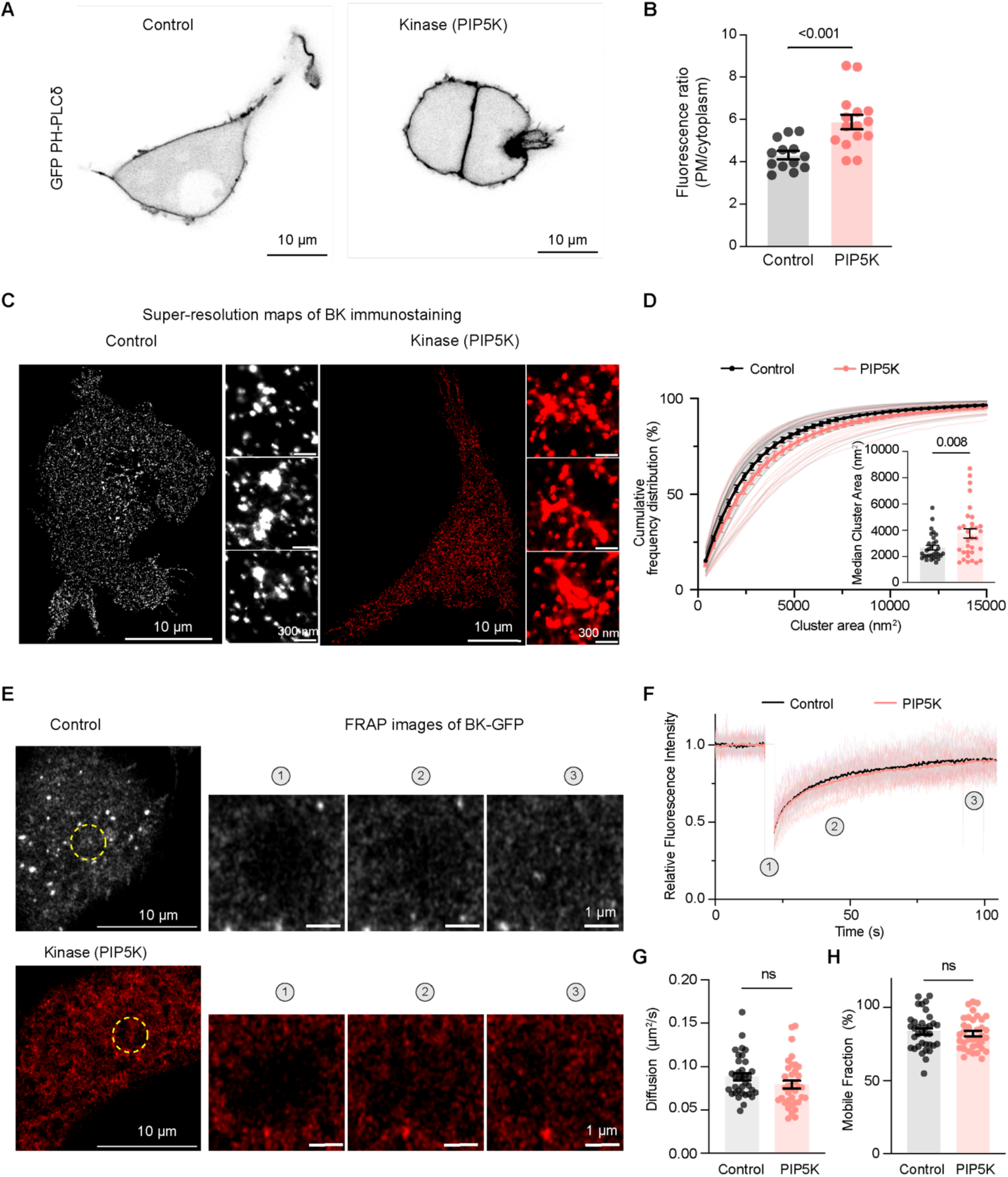
PtdIns(4,5)*P*_2_ enrichment promotes larger clusters without affecting BK channel mobility. **A.** Representative confocal images of tsA-201 cells transfected with PH-PLCδ. Right: control cells; left: cells transfected with PIP5K. Scale bar: 10 µm. **B.** Quantification of plasma membrane to cytosol PH-PLCδ intensity ratio between control (n = 13) and PIP5K expressing cells (n = 15). **C.** Representative super-resolution localization maps of BK immunostaining [use this word in figure too]. Right: control cells; left: cells transfected with PIP5K. Scale bars: 10 µm and 300 nm in magnification. **D.** Cumulative frequency plot comparing BK cluster size distributions between control (n = 31) and PIP5K (n = 32) overexpressing cells. Statistical significance was tested using a 2-way ANOVA. P-value was 0.001. Inset: scatter plot showing median cluster size (nm²). Statistical significance was tested using a two-tail t-test. **E.** Representative FRAP time-lapse images of BK-GFP top: control, bottom: cells transfected with PIP5K. Scale bars: 10 µm and 1 µm in magnification **F.** FRAP recovery curves comparing BK-GFP mobility between control and cells transfected with PIP5K. **G.** Scatter dot plot comparing BK clusters diffusion between control (n = 35) and cells transfected with PIP5K (n = 36). **H**. Scatter dot plot comparing mobile fraction of BK clusters in control and cells transfected with PIP5K. Statistical significance in all scatter plots was tested using a two-tail t-test.

Next, to determine whether PtdIns(4,5)*P*_2_ enrichment influences BK channel mobility, we performed FRAP analysis on BK-GFP–expressing cells (Figure 2E). Fluorescence recovery curves revealed comparable kinetics between control and PIP5K-expressing cells (Figure 2F), and quantification of diffusion coefficients showed no significant difference (control: 0.088 ± 0.004 μm²/s; PIP5K: 0.080 ± 0.004 μm²/s; Figure 2G), indicating that PtdIns(4,5)*P*_2_ enrichment does not alter the lateral diffusion of BK channels. Consistent with this, the mobile fraction of BK channels remained unchanged from 84 ± 2% in control to 82 ± 2% in PIP5K-expressing cells, indicating that elevated PtdIns(4,5)P_2_ levels do not alter the lateral mobility of BK channels.

After establishing that PtdIns(4,5)P_2_ enrichment alters BK clustering without affecting channel mobility, we next examined how PtdIns(4,5)P_2_ depletion influences BK organization and dynamics. Chronic depletion was achieved by overexpressing PIP5P, which converts PtdIns(4,5)P_2_ to PI(4)P (Figure S2-1). Using the PH-PLCδ-GFP biosensor, we confirmed that PIP5P expression reduced plasma membrane PtdIns(4,5)P_2_ levels by around 18% (Figure 3A-B).

**Figure 3.**
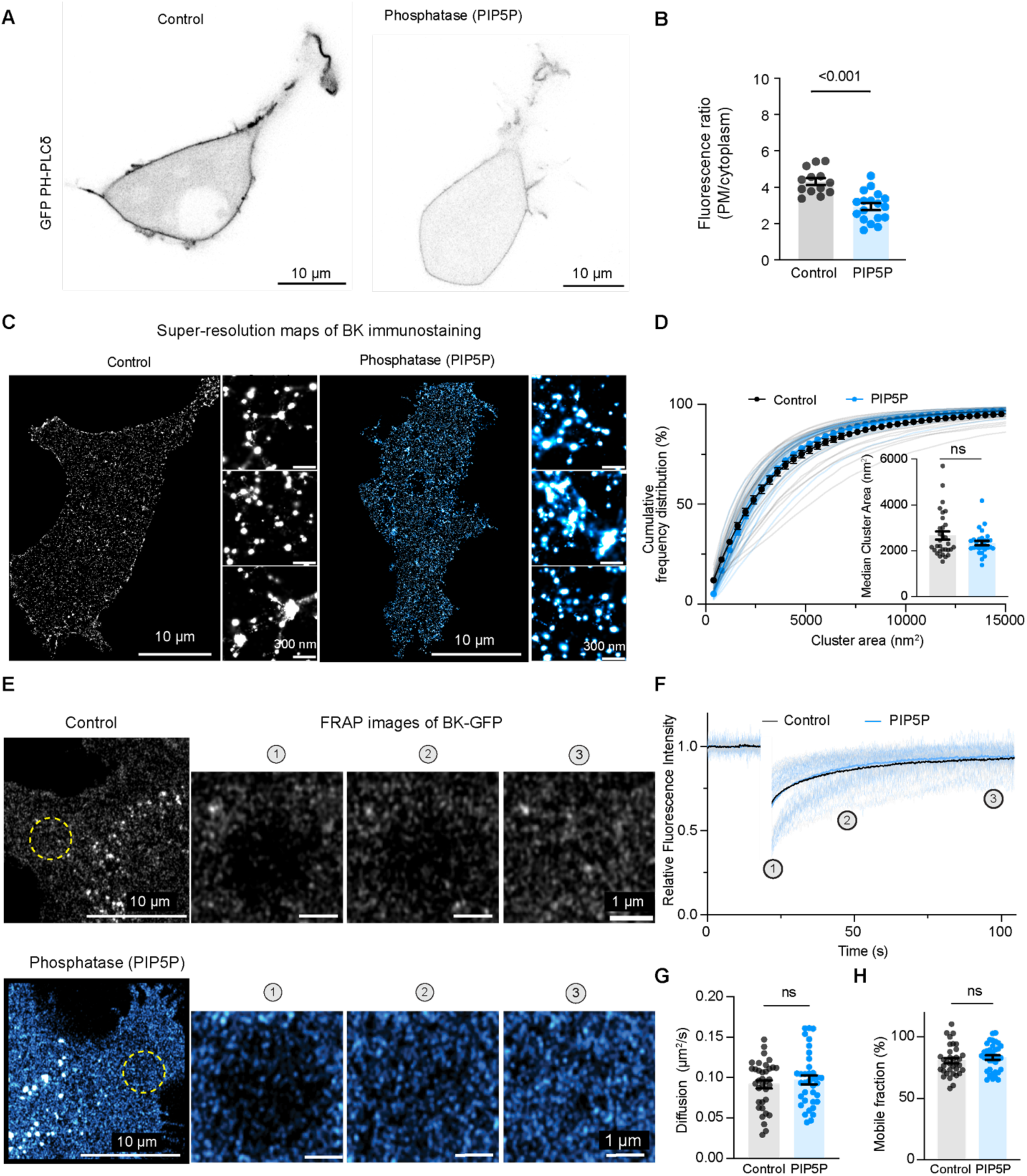
Long-term PtdIns(4,5)*P*_2_ depletion does not affect BK organization or mobility. **A.** Representative confocal images of tsA-201 cells transfected with PH-PLCδ. Right: control cells; left: cells transfected with PIP5P. Scale bar: 10 µm. **B.** Quantification of plasma membrane to cytosol PH-PLCδ intensity ratio between control (n = 13) and PIP5P expressing (n = 15) cells. Statistical significance was tested using a two-tail t-test. **C**. Representative super-resolution localization maps of BK immunostaining. Right: control cells; Left: cells transfected with PIP5P. Scale bars: 10 µm and 300 nm in magnification. **D.** Cumulative frequency plot comparing BK cluster size distributions between control (n = 31) and PIP5P expressing (n = 29) cells. Inset: scatter plot showing median cluster size (nm²). Statistical significance was tested using a two-tail t-test. **E.** Representative FRAP time-lapse images of BK-GFP top: control; bottom: cells transfected with PIP5P: Scale bar: 10 µm and 1 µm in magnification **F.** FRAP recovery curves comparing BK-GFP mobility between control and cells transfected with PIP5P. **G.** Scatter dot plot comparing BK clusters diffusion between control (n= 35), cells transfected with PIP5P (n= 36) **H**. Scatter dot plot comparing mobile fraction of BK clusters in control and cells transfected with PIP5P. Statistical significance was tested using a two-tail t-test.

Chronic PtdIns(4,5)P_2_ depletion via PIP5P expression left BK cluster architecture unchanged (Figure 3C-D). Cluster size distributions overlapped between control and PIP5P-expressing cells (p-value = 0.2); and the median cluster size did not change (2801 ± 147 nm² in control cells and 2334 ± 98 nm², p = 0.5). FRAP measurements revealed no significant differences in diffusion coefficients (control: 0.092 ± 0.005 μm^2^/s; PIP5P: 0.097 ± 0.005 μm^2^/s; Figure 3G) or mobile fractions (control: 81 ± 2 %; PIP5P: 83 ± 2 %; Figure 3H).

To evaluate the consequences of substantial and fast PtdIns(4,5)P_2_ loss on channel organization, we coexpressed BK and M1R and applied Oxo-M (Supplementary video S2-1). This treatment produced a pronounced reorganization of BK channels (Figure 4A). The cumulative frequency distribution was shifted toward larger cluster sizes (2-way ANOVA between cumulative curves, *p* = 0.006). Consistent with this shift, the median cluster size increased from 2071 ± 92 nm^2^ in control cells to 2728 ± 153 nm^2^ after Oxo-M treatment (*p* = 0.0005), representing a difference of 657 nm^2^ (Figure 4B).

**Figure 4.**
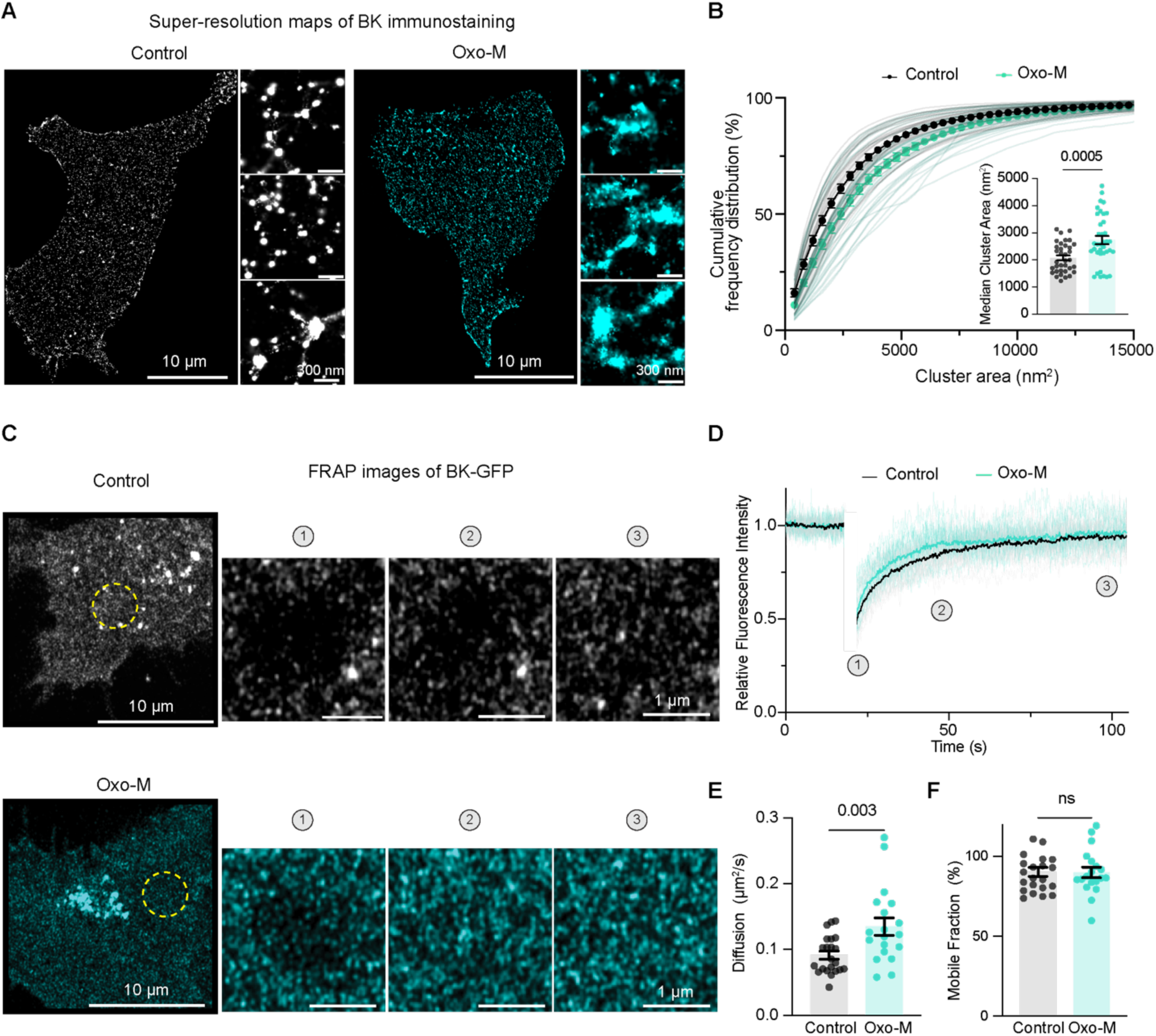
Substantial and fast PtdIns(4,5)P_2_ depletion promotes larger clusters and reduces BK mobility. **A**. Representative super-resolution localization maps of BK immunostaining. Right: control cells; Left: cells treated with Oxo-M. Scale bars: 10 µm and 300 nm in magnification. **B.** Cumulative frequency plot comparing BK cluster size distributions between control (n = 35) and Oxo-M-treated cells (n = 37). Inset: scatter plot showing median cluster size (nm²). Statistical significance was tested using a two-tail t-test. **C.** Representative FRAP time-lapse images of BK-GFP top: control; bottom: cells treated with Oxo-M: Scale bar: 10 µm and 1 µm in magnification. **D.** FRAP recovery curves comparing BK-GFP mobility between control and Oxo-M. **E.** Scatter dot plot comparing BK clusters diffusion between control (n = 22) and Oxo-M (n = 19). **F**. Scatter dot plot comparing mobile fraction of BK clusters in control and cells transfected with PIP5P. Statistical significance was tested using a two-tail t-test.

Next, we performed FRAP on cells coexpressing BK-GFP and M1R. Since PtdIns(4,5)P_2_ resynthesis limits the duration of M1R-dependent depletion, Oxo-M treatment was combined with GSK-A1, a PI4KIIIα inhibitor that prevents replenishment of the PtdIns(4,5)P_2_ pool. Under these conditions, BK channels exhibited faster fluorescence recovery (Figure 4C-D), with diffusion coefficients increasing from 0.091 ± 0.006 μm^2^/s in control cells to 0.135 ± 0.013 μm^2^/s following combined Oxo-M and GSK-A1 treatment (Figure 4E). The mobile fraction remained unchanged (control: 90 ± 3%; Oxo-M: 90 ± 3%; Figure 4F), indicating that acute PtdIns(4,5)P_2_ depletion increases BK diffusion rates without altering the proportion of mobile channels.

### Actin disruption reduces BK cluster size without altering channel mobility

To further investigate how membrane architecture shapes BK channel organization, we examined the contribution of the cortical actin cytoskeleton. We disrupted filamentous actin using cytochalasin D (Cyt D), a pharmacological inhibitor of actin polymerization, and visualized actin integrity using Actin CellMask. Compared to control cells, Cyt D treatment resulted in a marked reduction of cortical filamentous structures, confirming effective cytoskeletal disruption (Figure 5A-B). Next, we investigated how BK channel clustering is influenced by the integrity of the actin cytoskeleton. Representative super-resolution maps of BK clusters under control and Cyt D treatment are shown in Figure 5C. Disruption of actin filaments shifted the cumulative frequency distribution toward smaller cluster sizes (Figure 5D; 2-way ANOVA between cumulative curves, *p* < 0.0001). Consistent with this shift, the median cluster size decreased from 2368 ± 122 nm² in control cells to 1547 ± 102 nm² after Cyt D treatment (*p* < 0.0001), representing a difference of 821 nm².

**Figure 5.**
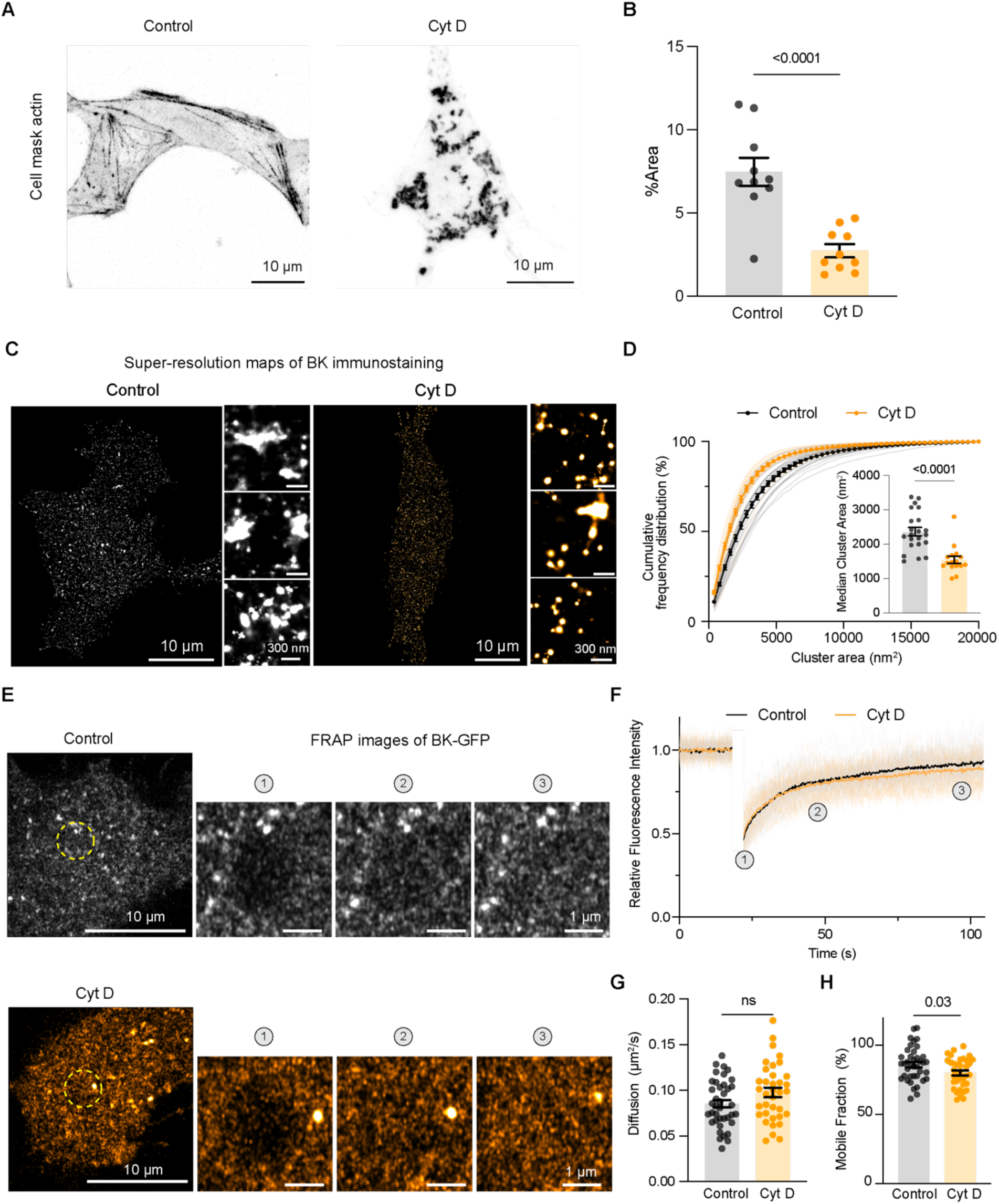
Actin disruption reduces BK cluster size without altering channel mobility. **A.** Representative confocal images of tsA-201 cells stained with actin cell mask. Right: control cells; left: cells treated with Cyt D. Scale bar: 10 µm. **B.** Quantification of fractional actin cell mask area (control n = 13; Cyt D n = 15). Statistical significance was tested using a two-tail t-test. **C.** Representative super-resolution localization maps of BK immunostaining. Right: control cells (n = 22); left: Cyt D treated (n = 15). Scale bars: 10 µm and 300 nm in magnification. **D.** Cumulative frequency plot comparing BK cluster size distributions between control and Cyt D-treated cells. Inset: scatter plot showing median cluster size in nm². Statistical significance was tested using a two-tailed t-test. **E.** Representative FRAP time-lapse images of BK-GFP, top: control, bottom: Cyt D treatment. Scale bars: 10 µm and 1 µm in magnification. **F.** FRAP recovery curves comparing BK-GFP mobility between treated (n = 36) and control (n = 39) cells. **G.** Scatter dot plot comparing BK channel diffusion between treated and control cells. **H.** Scatter dot plot comparing mobile fraction of BK clusters in treated and control cells. Statistical significance was tested using a two-tail t-test.

To evaluate whether actin disruption affects BK channel mobility, we performed FRAP following Cyt D treatment (Figure 5E-F). Recovery kinetics showed modest alterations, with no significant change in diffusion coefficient between conditions (control: 0.085 ± 0.004 μm^2^/s; Oxo-M: 0.098 ± 0.005 μm^2^/s; Figure 5G). The mobile fraction showed a modest decrease under Cyt D treatment from 86 ± 2% in control cells to 80 ± 2% with Cyt D treatment (Figure 5H).

### Endogenous BK channel clustering and density are regulated by cholesterol, PtdIns(4,5)P_2_, and actin cytoskeleton

After characterizing BK channel clustering in a heterologous expression system, we next sought to examine whether similar organizational principles apply in a physiologically relevant, endogenous context. To this end, we analyzed BK clusters in INS-1 cells, a pancreatic β-cell line that endogenously expresses BK channels. In this context, we also tested the effect of cholesterol, PtdIns(4,5)P_2_, and actin on BK channel expression density at the plasma membrane.

Cholesterol depletion with MβCD led to a marked reduction in BK cluster density, from 3.5 ± 0.1 clusters/μm² in control cells to 2.7 ± 0.1 clusters/μm² after treatment (Figure 6A–B). Cholesterol depletion also shifted the cumulative frequency distribution toward smaller cluster sizes (Figure 6C, 2-way ANOVA between cumulative curves, *p =* 0.0001). Consistent with this leftward shift, the median cluster size decreased from 3296 ± 142 nm² in control cells to 2684 ± 187 nm² after MβCD treatment (*p* = 0.01), representing a decrease of 612 nm², indicating that cholesterol also contributes to BK channel organization in INS-1 cells.

**Figure 6.**
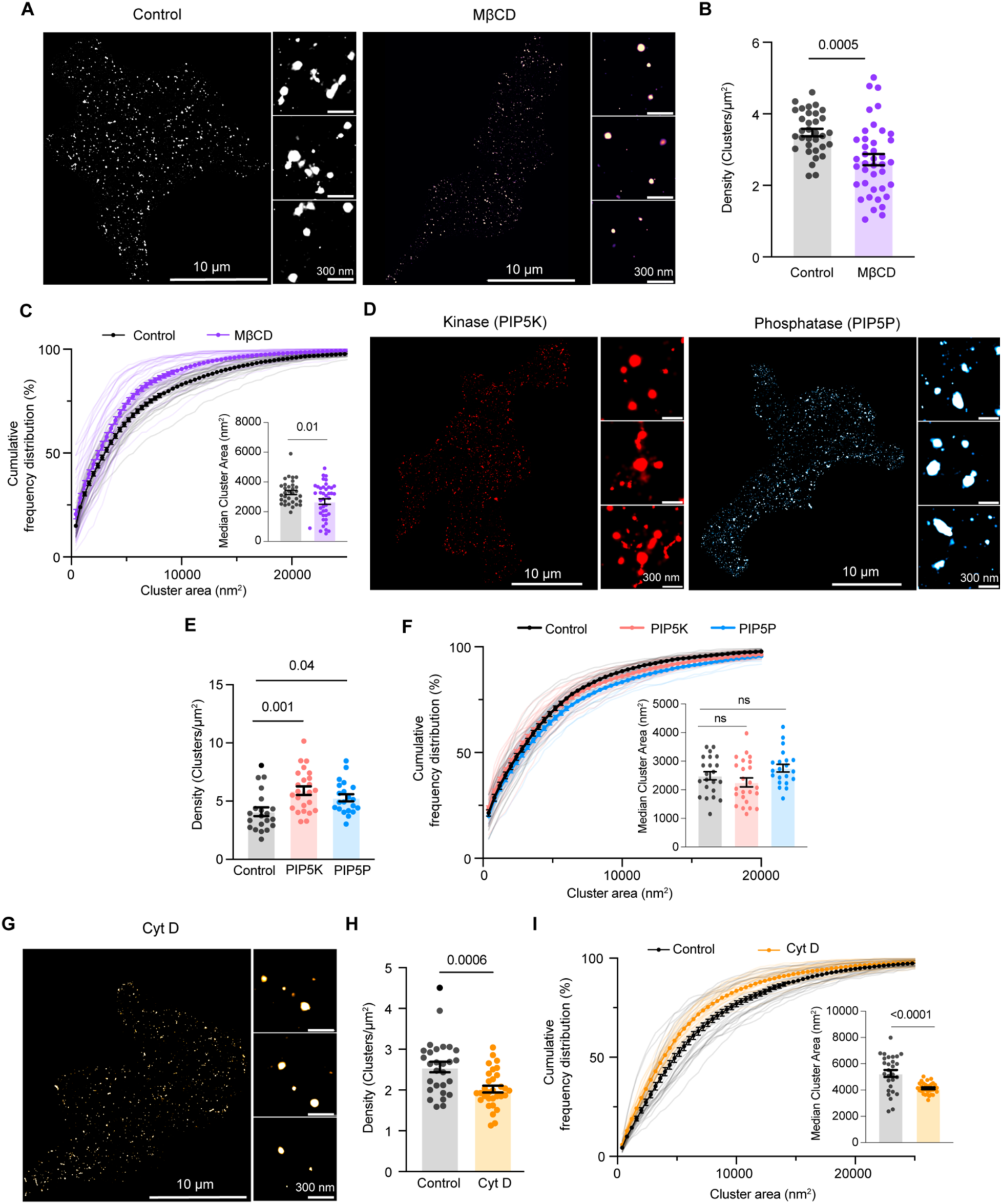
Endogenous BK channel clustering and density are regulated by cholesterol, PtdIns(4,5)P_2_, and the actin cytoskeleton. **A.** Representative super-resolution localization maps of BK immunostaining in INS-1 cells under endogenous expression. Right: control; left: MβCD-treated cells. Scale bars: 10 µm and 300 nm (inset). **B.** Scatter dot plot comparing BK cluster density between control and MβCD-treated INS-1 cells. **C.** Cumulative frequency plot of BK cluster size distributions in control versus MβCD-treated cells. Inset: scatter plot showing median cluster size (nm²). **D.** Representative super-resolution localization maps of BK immunostaining in INS-1 cells expressing right: PIP5K, left: PIP5P. Scale bars: 10 µm and 300 nm (inset). **E.** Scatter dot plot comparing BK cluster density between control, PIP5K-expressing, and PIP5P-expressing INS-1 cells. **F.** Cumulative frequency plot of BK cluster size distributions in control versus PIP5K-expressing, and PIP5P-expressing INS-1 cells. Inset: scatter plot showing median cluster size (nm²). **G.** Representative super-resolution localization maps of BK immunostaining in INS-1 cells treated with cytochalasin D (Cyt D). Scale bars: 10 µm and 300 nm (inset). **H.** Scatter dot plot comparing BK cluster density between control and Cyt D-treated INS-1 cells. **I.** Cumulative frequency plot of BK cluster size distributions in control versus Cyt D-treated cells. Inset: scatter plot showing median cluster size (nm²). Statistical significance was tested using a two-tail t-test.

Manipulation of PtdIns(4,5)P_2_ levels through PIP5K overexpression produced a significant increase in BK cluster density relative to control, rising from 4.1 ± 0.4 to 5.9 ± 0.3 clusters/μm². A similar, though more modest, increase was observed in PIP5P-expressing cells (5.3 ± 0.3 clusters/μm²; control: 4.1 ± 0.4 clusters/μm²; Figure 6D–E). PtdIns(4,5)P_2_ enrichment did not shift the cumulative frequency distribution (Figure 6F, 2-way ANOVA between cumulative curves*, p =* 0.5), while PtdIns(4,5)P_2_ depletion shifted the cumulative frequency distribution toward larger cluster sizes (Figure 6F, 2-way ANOVA between cumulative curves, *p =* 0.001). However, the change in cluster size was relatively small, resulting in no significant difference between the medians (control: 2533 ± 143 nm²; PIP5K: 2257 ± 155 nm²; PIP5P: 2757 ± 134 nm²).

Disruption of the actin cytoskeleton with Cyt D significantly reduced BK cluster density, decreasing from 2.6 ± 0.1 clusters/μm² in control cells to 2.0 ± 0.1 clusters/μm² (Figure 6G–H). Disruption of actin filaments shifted the cumulative frequency distribution toward smaller cluster sizes (Figure 6I; 2-way ANOVA between cumulative curves, *p* = 0.0003). Consistent with this shift, the median cluster size decreased from 5263 ± 265 nm² in control cells to 4133 ± 73 nm² after Cyt D treatment (*p* < 0.0001), representing a difference of 1130 nm².

### Density, but not diffusion, regulates BK channel clustering

To evaluate how different experimental treatments influence the nanoscale organization of BK channels, we examined the relationship between treatment-induced changes in cluster size and two key parameters: diffusion and cluster density. In tsA-201 cells, relative median cluster area (treatment/control) showed no meaningful association with relative diffusion across treatments (Figure 7A; R^2^ = 0.02), indicating that changes in lateral mobility do not correspond to alterations in cluster size in this overexpression system. Because cluster density cannot be meaningfully interpreted in an overexpression context, we assessed the relationship between density and cluster size in INS-1 cells, where channels are expressed endogenously. Here, treatments produced a strong positive correlation (Figure 7B; R^2^ = 0.96) between relative cluster density and relative median cluster area, demonstrating that conditions that increase the number of clusters per unit area also proportionally increase cluster size.

**Figure 7.**
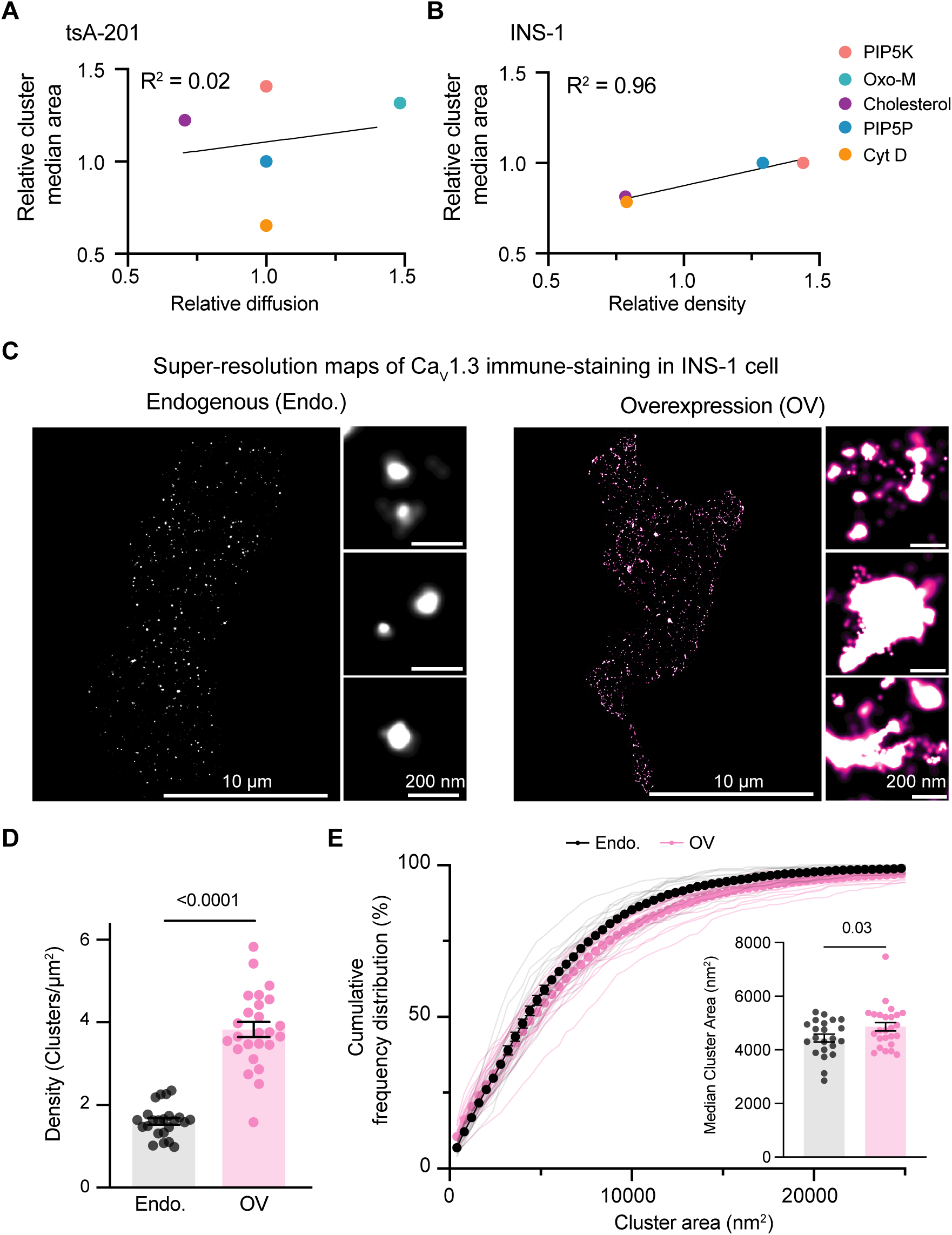
Density but not diffusion regulates BK channel clustering. **A.** Relationship between treatment-induced changes in BK channel cluster size and relative diffusion in tsA-201 cells. **B.** Relationship between treatment-induced changes in BK channel cluster size and cluster density in INS-1 cells. **C.** Representative super-resolution localization maps of Ca_V_1.3 immunostaining in INS-1 cells. Left: endogenous, right: overexpression. **D.** Scatter dot plot comparing Ca_V_1.3 cluster density between endogenous (n = 22) and overexpression (n = 25). **E.** Cumulative frequency plot of Ca_V_1.3 cluster size distributions in endogenous versus overexpression. Inset: scatter plot showing median cluster size (nm²). Statistical significance was tested using two-way ANOVA across cumulative curves and a two-tail t-test.

Based on our model of BK channel organization, one prediction is that overexpressing an endogenously expressed channel—or more generally, increasing the surface density of any ion channel—should lead to a corresponding increase in cluster size. To test this prediction, we used INS-1 cells and overexpressed the voltage-gated calcium channel Ca_V_1.3. Representative dSTORM maps are shown in Figure 7C. As expected, overexpression produced a substantial increase in channel density (Figure 7D; *p* < 0.0001). In support of our model, the cumulative frequency distribution of cluster area shifted toward larger values in the overexpression condition (Figure 7E; two-way ANOVA across cumulative curves, p < 0.0001). Consistent with this shift, the median cluster area increased from 4443 ± 145 nm² under endogenous expression to 4861 ± 158 nm² following overexpression (p = *0.03*), representing an increase of 420 nm².

## Discussion

Many transmembrane proteins, including BK channels, assemble into membrane clusters [16, 17]. Despite the ubiquity of cluster formation, there is no consensus on a generalized mechanism underlying this phenomenon. Several “attractor mechanisms” have been proposed—scaffolding interactions [18], lipid rafts [19, 20], and membrane curvature [21, 22]—all of which can recruit proteins into discrete membrane domains. Whether clustering arises primarily from diffusion-driven encounters [23] or from insertion into pre-existing nucleation sites [24] also remains debated. By perturbing cholesterol, PtdIns(4,5)P_2_, and the actin cytoskeleton, our work provides new insight into which of these mechanisms may be most relevant for BK channels.

BK channels are enriched in cholesterol-rich, detergent-resistant membranes [25, 26] and directly bind cholesterol via CRAC motifs [10]. Consistent with this, cholesterol depletion reduced both BK diffusion and mobile fraction, in agreement with single-particle tracking studies showing near-immobilization after MβCD treatment [27]. Cholesterol depletion also altered BK cluster size, but in a cell-type-specific manner: cluster area increased in tsA-201 cells yet decreased in INS-1 cells. Similar bidirectional effects have been reported for other channels and transporters, including the dopamine transporter [3] and Kv2.1 [28]. The literature has long emphasized that cholesterol regulates BK channels in a highly context-dependent manner, often attributed to auxiliary β subunits [29]. This aligns with our observation that INS-1 cells, which express β subunits, respond oppositely to tsA-201 cells, which do not express β subunits. Although no single pattern emerges across cell types, our findings reinforce the view that cholesterol modulates BK nanoscale organization in a manner shaped by cellular context.

PtdIns(4,5)P_2_ also influenced BK mobility and clustering but with distinct outcomes depending on the mode of perturbation. Although direct interactions between PtdIns(4,5)P_2_ and basic residues in the S6–S7 linker enhance BK activity [30], chronic enrichment or depletion did not alter diffusion. In contrast, rapid and substantial PtdIns(4,5)P_2_ depletion increased BK mobility, consistent with simulations of Kir2.2 channels [31] and theoretical predictions that reduced PtdIns(4,5)P_2_ should enhance lateral diffusion [32]. Similar lipid-dependent effects on clustering have been observed in other systems, such as αvβ3 integrins following neomycin-mediated sequestration [33]. Beyond mobility, PtdIns(4,5)P_2_ perturbations reorganized BK channels at the nanoscale: both enrichment and rapid depletion shifted channels toward larger clusters, and PIP5K-mediated enrichment increased cluster density in INS-1 cells. These results suggest that PtdIns(4,5)P_2_ acts not as a linear regulator but as a homeostatic organizer of BK architecture, where deviations in either direction promote the formation of larger nanodomains.

The actin cytoskeleton contributed in yet another way. BK channels interact with actin through cortactin [9], leading us to expect that actin disruption would increase mobility. Surprisingly, cytochalasin D did not alter diffusion and produced only a modest reduction in mobile fraction. This contrasts with prior work showing increased BK diffusion after latrunculin A treatment [34], a discrepancy that may reflect drug-specific effects, as latrunculin A and cytochalasin D differentially influence the mobility of other membrane proteins [35]. In contrast to its minimal effect on mobility, actin disruption markedly reduced BK cluster size in both cell types, indicating a conserved role for actin in maintaining BK nanodomains. Actin perturbation has been shown to either increase or decrease cluster size depending on the protein—examples include Glut1 [36], E-cadherin [37], and Kv2.1 [28]—highlighting that membrane proteins engage with the cytoskeleton through diverse mechanisms. Collectively, these findings support a broad role for actin in stabilizing nanoscale protein organization.

Super-resolution imaging enabled us to quantify both cluster size and density in endogenous systems. Treatment-induced changes in these parameters were positively correlated, consistent with observations in Cav1.2 channels [24] and simulations of Kir2.2 [31]. These results suggest that BK clustering is strongly influenced by channel density. Notably, we found no relationship between changes in diffusion and cluster size, challenging the common assumption that clustering is diffusion-limited [38] and contrasting with recent proposals that BK clusters arise through diffusion-driven encounters [23, 39]. Instead, our data indicate that BK clustering is independent of diffusion.

## Conclusion and future directions

Across the perturbations tested, actin emerged as the strongest determinant of BK cluster size, with depolymerization producing smaller clusters in both cell types. By comparing cholesterol, PtdIns(4,5)P_2_, and actin perturbations, we also identified density—but not diffusion—as a key correlate of BK cluster size, challenging diffusion-driven models of BK organization. Future work should dissect how actin contributes to BK clustering, whether through physical corralling [40], interactions with scaffolding proteins [41], or both.

## Acknowledgement

We thank the BRIGHT-UP high-school student Karla Osnaya, and Roxanne Madden for their technical support. We thank Dr. Bertil Hille and Wendy Piñon-Teal for their critical reading of the manuscript. This study was supported by the National Institutes of Health MIRA R35 GM142690 to O.V. and the NHLBI R01 HL162609 to C.M. C.M is a Freeman Hrabowski HHMI Scholar.

---

**Figure S1-1.**
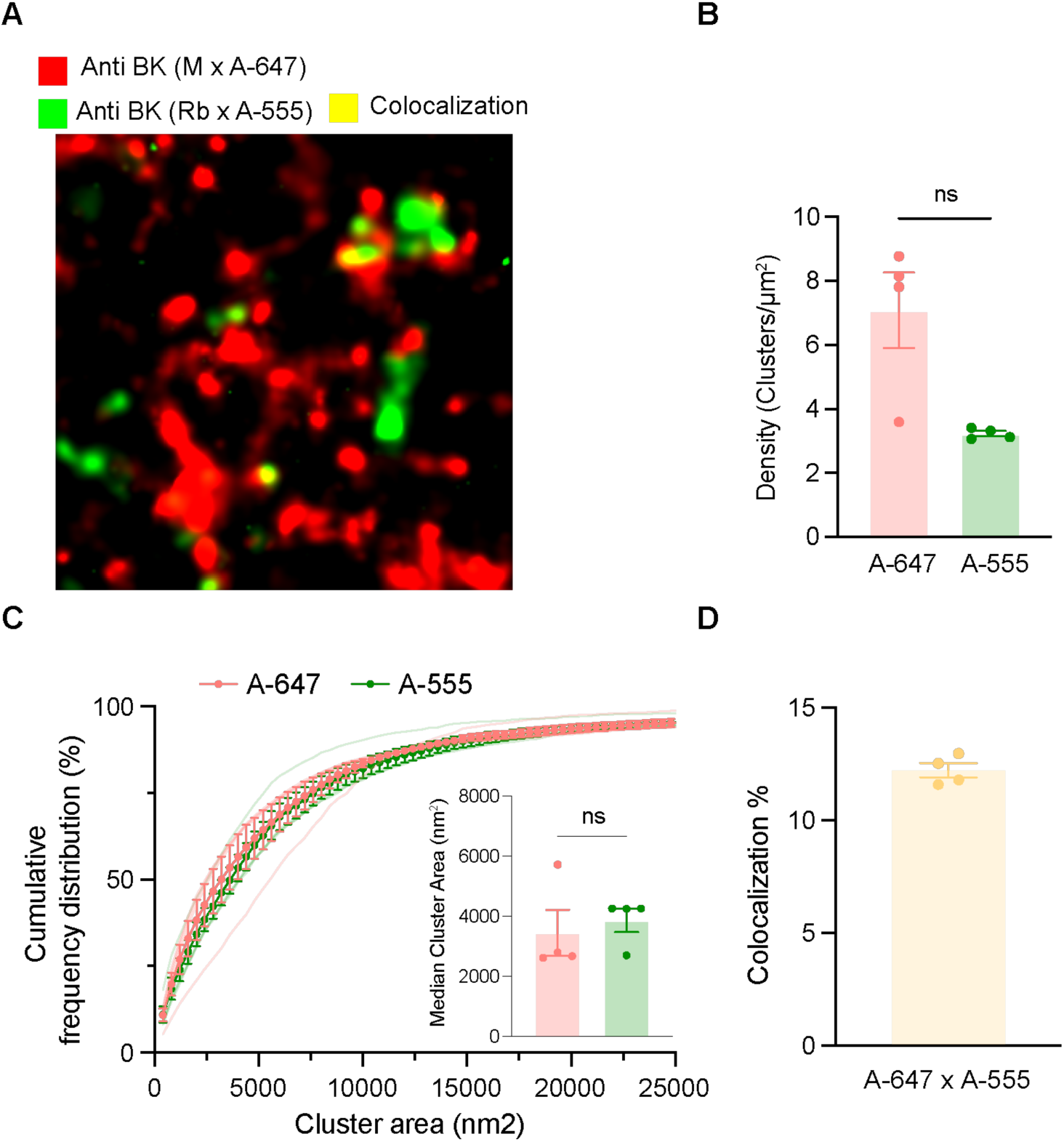
Limited antibody colocalization supports single-site binding per BK tetramer. **A.** Representative colocalization maps of the BK channel probed with two primary antibodies raised in mouse (red, Alexa-647) and rabbit (green, Alexa-555). **B.** Scatter plot showing BK cluster density (Clusters/μm^2^) **C.** Relative cumulative frequency distribution of BK channels probed with two primary antibodies raised in mouse (red, Alexa-647) and rabbit (green, Alexa-555). Inset: scatter plot showing median cluster size (nm²). **D.** Scatter plot showing colocalization between Alexa-647 and Alexa-555 signals. Data are from 4 cells shown as mean ± SEM.

**Figure S1-2.**
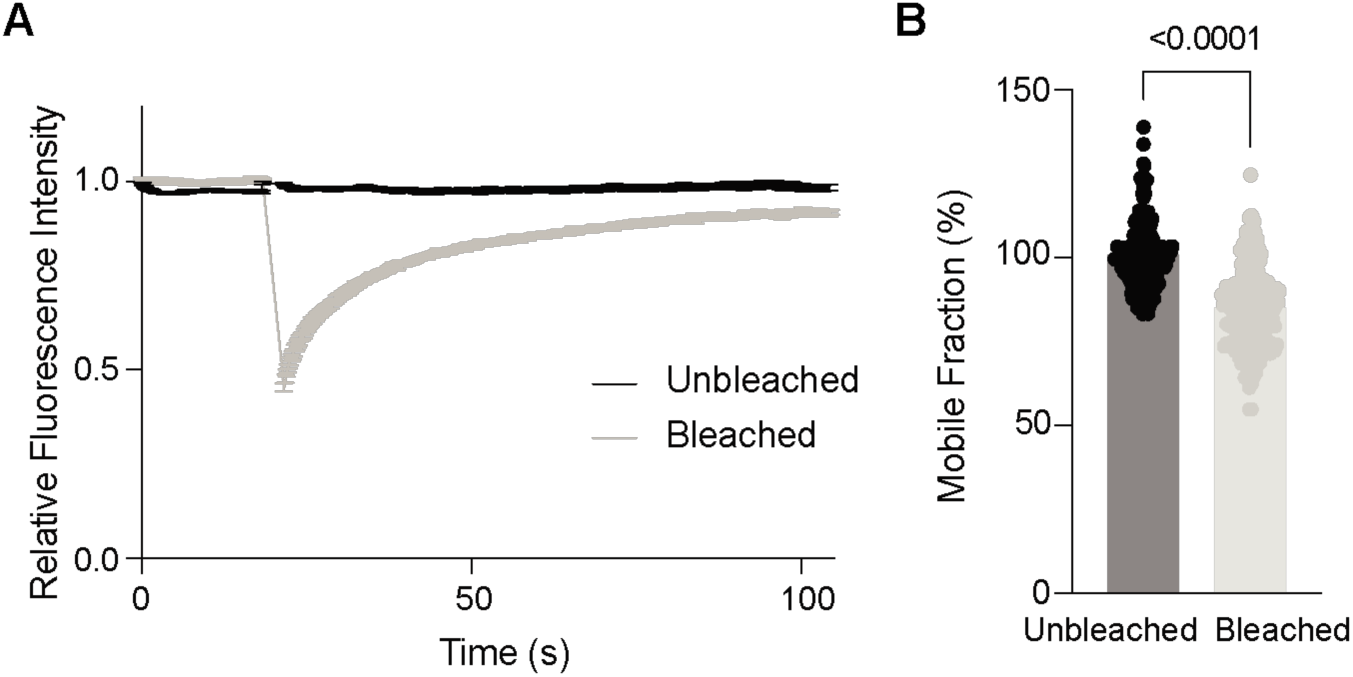
Comparison of the change in fluorescence between bleached and unbleached areas. **A**. Time course of fluorescence intensity from bleached (grey) and unbleaced (black) areas in tsA-201 cells transfected with fluorescently-tagged BK channels. **B**. Comparison of the percentage of change between the initial and final fluorescence fraction, representing the percentage mobile fraction. Statistical significance was tested using a two-tail t-test.

**Figure S2-1.**
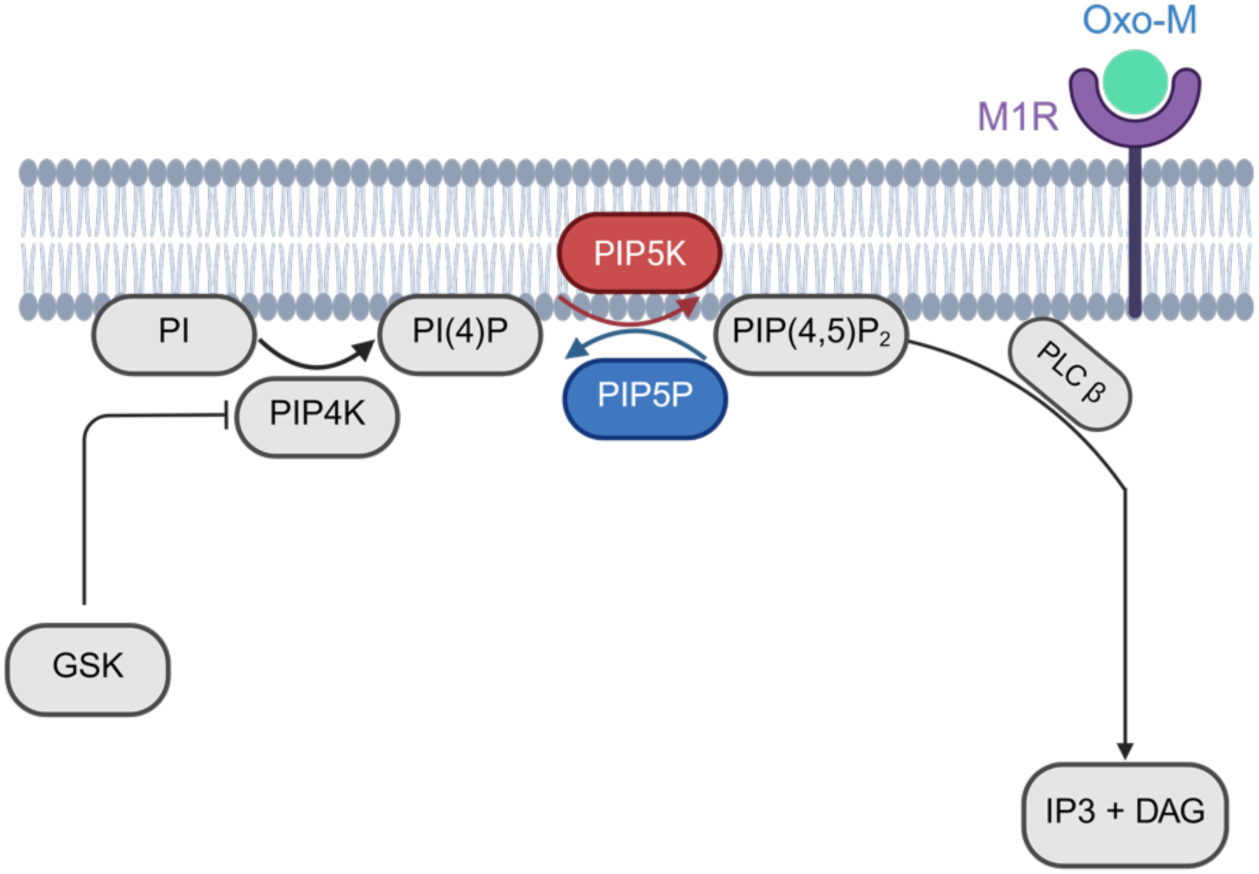
Schematic illustrating experimental strategies used to manipulate PtdIns(4,5)P_2_ abundance. Chronic modulation was achieved by overexpressing PIP5K or PIP5P to increase or decrease PtdIns(4,5)P_2_ synthesis, respectively. Acute depletion was induced by activating the M1 muscarinic receptor (M1R) with Oxotremorine-M (Oxo-M), which stimulates PLCβ-mediated hydrolysis of PtdIns(4,5)P_2_, and inhibiting the PtdIns(4,5)*P*_2_ synthesis pathway with GSK.

**Supplementary video S2-1. Validation of approach for rapid and substantial decrease of PtdIns(4,5)*P*_2_.** Cell expressing the biosensor PH_PLCδ1_ to detect changes in PtdIns(4,5)*P*_2_ and the muscarinic acetylcholine receptor type 1 was stimulated with oxotremorine-M (Oxo-M) in the presence of the PI4K inhibitor GSK-A1. The levels of PtdIns(4,5)*P*_2_ were suppressed for 46 minutes as suggested by the increase in fluorescence in the cytosol.

